# Functionally asymmetric motor neurons coordinate locomotion of *Caenorhabditis elegans*

**DOI:** 10.1101/244434

**Authors:** Oleg Tolstenkov, Petrus Van der Auwera, Jana F. Liewald, Wagner Steuer Costa, Olga Bazhanova, Tim Gemeinhard, Amelie C. F. Bergs, Alexander Gottschalk

## Abstract

Invertebrate nervous systems are valuable models for fundamental principles of the control of behavior. Ventral nerve cord (VNC) motor neurons in *Caenorhabditis elegans* represent one of the best studied locomotor circuits, with known connectivity and functional information about most of the involved neuron classes. However, for one of those, the AS motor neurons (AS MNs), no physiological data is available. Combining specific expression and selective illumination, we precisely targeted AS MNs by optogenetics and addressed their role in the locomotion circuit. After photostimulation, AS MNs induce currents in post-synaptic body wall muscles (BWMs), exhibiting an initial asymmetry of excitatory output. This may facilitate complex regulatory motifs for adjusting direction during navigation. By behavioral and photo-inhibition experiments, we show that AS MNs contribute to propagation of the antero-posterior body wave during locomotion. By Ca^2+^-imaging in AS MNs and in their synaptic partners, we also reveal that AS MNs play a role in mediating forward and backward locomotion by integrating activity of premotor interneurons (PINs), as well as in coordination of the dorso-ventral body wave. AS MNs do not exhibit pacemaker properties, but potentially gate VNC central pattern generators (CPGs), as indicated by ceasing of locomotion when AS MNs are hyperpolarized. AS MNs provide positive feedback to the PIN AVA via gap junctions, a feature found also in other locomotion circuits. In sum, AS MNs have essential roles in coordinating locomotion, combining several functions, and emphasizing the compressed nature of the *C. elegans* nervous system in comparison to higher animals.

**Highlights:** A class of motor neurons with unidentified function – AS cholinergic motor neurons - was characterized in *C. elegans*.

AS neurons show asymmetry in both input and output and are specialized in coordination of dorso-ventral undulation bends.

AS neurons mediate antero-posterior propagation of the undulatory body wave during locomotion.

AS neurons integrate signals for forward and reverse locomotion from premotor interneurons and may gate ventral nerve cord central pattern generators (CPGs) via gap junctions.

## Introduction

Locomotion represents a basic component of many complex behaviors and is regulated by neuronal circuits that share similar properties in a wide variety of species, including humans (Guertin, 2013; Kiehn, 2011; Mullins et al., 2011). These circuits, as shown in virtually all model systems studied, can generate rhythmic motor patterns without sensory inputs, and therefore act as CPGs (Pearson, 1993). In higher vertebrates CPGs are very complex systems and represent distributed networks made of multiple coupled oscillatory centers, grouping in pools as discrete operational units (Fidelin et al., 2015; Kiehn et al., 2010; Rybak et al., 2015). Motor neurons (MNs) within the pool are able to integrate convergent inputs; they are recruited and activated gradually, which underlies the variable changes in muscle tension that are necessary for movement. Overlaid on these circuits are interactions with (and between) premotor interneurons (PINs, also called command interneurons), which modulate the patterns of MN activity and coordinate the CPGs (Goulding, 2009). In mammals, it is particularly the commissural interneurons (CIN) which regulate activity of left and right CPGs, and which may therefore themselves act as rhythm generators. Such neurons are often excitatory (e.g. CINei of the mouse; Kiehn, 2016), but can also be inhibitory (e.g. CINi), while others act as electrical connectors or activity ‘sinks’ for CPGs and motor neuron pools. An example are ipsilateral V2a interneurons in zebrafish, which are retrogradely recruited by motor neurons (Song et al., 2016) to modulate their activity.

Despite the difference in the forms of locomotion and anatomy of neural circuits between vertebrates and invertebrates, they share similar principles. Yet, how complex vertebrate locomotion circuits operate and how they developed from more simple ones is not understood in its entirety, thus a comprehensive analysis of invertebrate circuits is a prerequisite to this goal. The relative simplicity of invertebrate nervous systems has helped to develop concepts that guide our understanding of how complex neuronal networks operate (Marder et al., 2005; Selverston, 2010). *C. elegans* is a nematode with only 302 neurons in the hermaphrodite. A fully reconstructed wiring diagram of its neural circuits (Varshney et al., 2011; White et al., 1986) and various tools for imaging and (opto)genetic interrogation of circuit activity (Fang-Yen et al., 2015; Leifer et al., 2011; Nagel et al., 2005; Stirman et al., 2011) render *C. elegans* a useful model to study fundamental principles of the neuronal control of behavior.

*C. elegans* moves by generating waves of dorso-ventral bends along its body. These predominantly lead to forward movement, which is occasionally interrupted by brief backing episodes, the frequency of which is modulated by sensory responses (Cohen and Sanders, 2014; Gjorgjieva et al., 2014; Pierce-Shimomura et al., 2008; Zhen and Samuel, 2015). The animal’s undulations are controlled by neural circuits in the head and VNC. The core components of the motor circuits in *C. elegans* include head motor/interneurons that exhibit oscillations during alternating head bending (Hendricks et al., 2012; Shen et al., 2016), and which are transmitted to the remainder of the body (mainly) by proprioceptive feedback (Wen et al., 2012). In the body, motor neurons are found in ensembles or subcircuits, repeating 6 times from the ‘neck’ to the tail of the animal, containing one or two neurons of each class (6–13 neurons found in the individual classes, with 11 AS MNs; Haspel and O’Donovan, 2011; White et al., 1986). Upstream of the motor neurons are PINs which integrate inputs from sensory and other interneurons, and that relay their activity in a gating fashion: They are themselves not oscillatory, but set up- or down-states of the motor neurons, using gap junction networks, in a manner similar to the V2a interneurons of the fish (Song et al., 2016). The classes of MNs are distinguished by transmitter used (acetylcholine or GABA), ventral or dorsal innervation, and roles in forward or backward locomotion (Von Stetina et al., 2005; Zhen and Samuel, 2015). Functions of the different types of MNs are understood to various degrees. For example, the DA9 A-type MN was recently demonstrated to generate intrinsic rhythmic activity by P/Q/N-type Ca^2+^ channels, which is potentiated by activity of the reversal PIN AVA (Gao et al., 2017). Thus, motor neurons, rather than interneurons, can be oscillators, demonstrating that different activities are compressed in the *C. elegans* motor circuit with its limited number of cells. To fully understand these circuits, all of the motor neurons need to be characterized. However, for the cholinergic AS MN class, representing one fifth of VNC cholinergic neurons, surprisingly no physiological data is available. Yet, these neurons are interesting in that they asymmetrically innervate only dorsal muscle and ventral inhibitory VD neurons. Further, in contrast to other MN types, the AS MNs are innervated extensively by chemical synapses from both forward and reverse PINs, and they also form gap junctions with these cells.

In this study, we investigated the role of AS MNs in the VNC locomotor circuit based on predictions made from the wiring diagram, using optogenetic tools, electrophysiology, behavioral analysis, and Ca^2+^ imaging in immobilized and moving animals. We reveal important roles of AS MNs in dorso-ventral and antero-posterior coordination of undulations during locomotion, as stimulation of AS MNs distorts, and inhibition blocks, propagation of the body wave. We show that AS MNs act through excitation of dorsal muscles and inhibitory ventral VD motor neurons. The intrinsic activity of the AS MNs correlates best with forward locomotion, which corresponds to a stronger response of AS MNs to photodepolarization of the forward PIN AVB. Functionally asymmetric electrical connections suggest AS MN feedback control of the backward PIN AVA, a feature recently observed for locomotor circuits also in other animals (Matsunaga et al., 2017; Song et al., 2016).

## RESULTS

### Selective expression and activation of optogenetic tools in AS MNs

Six classes of cholinergic MNs are involved in mediating the dorso-ventral sinusoidal wave observed during locomotion of *C. elegans*: DA, VA, DB, VB, VC and AS. Up to date, no promotor exclusively triggering expression in AS MNs is known. To achieve specific activation of AS MNs, we used a subtractive approach for expression combined with selective illumination. The p*unc-17* promoter (*unc-17* encodes the vesicular acetylcholine transporter) drives expression in all cholinergic neurons including the MNs in the VNC. In combination with p*acr-5* (driving expression in the DB and VB MNs) and p*unc-4* (driving expression in the DA, VA and VC MNs), we could restrict expression of optogenetic tools to the AS MNs (**Fig. 1A, B**): Briefly, broad expression from p*unc-17* was suppressed in the DB, VB, DA, VA and VC neurons by expressing dsRNA constructs targeting the optogenetic tool using p*acr-5* and p*unc-4* promoters (**Fig. 1AI**). Alternatively, we used the Q system (Wei et al., 2012): We placed the QF transcriptional activator under the p*unc-17* promoter, thus driving expression of the optogenetic tool from constructs harboring the QUAS QF binding motif. To restrict expression to AS MNs in the VNC, we additionally used the QS suppressor under the p*acr-5* and p*unc-4* promoters (**Fig. 1AII**). Last, since these approaches still led to expression in additional cholinergic neurons in head and tail ganglia, we avoided activation of optogenetic tools in those cells by selective illumination of segments of the animals body that correspond to AS MNs (Husson et al., 2012; Stirman et al., 2011, 2012; **Fig. 1AIII**).

**Figure 1:**
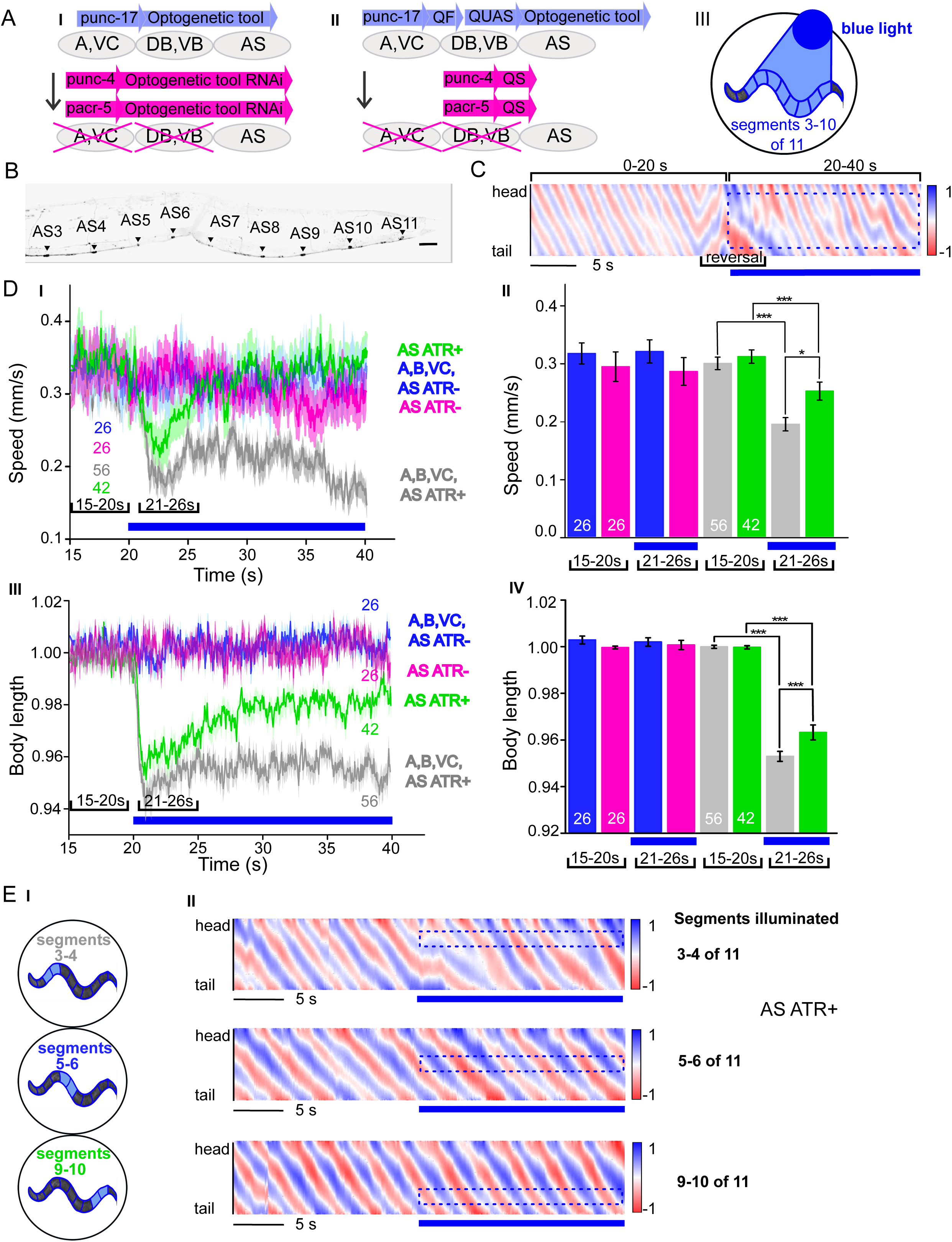
Specific photodepolarization of AS MNs via ChR2 leads to body contraction, increased bending angles and reduced speed in freely moving *C. elegans*: **A**) ‘Subtractive’ expression and illumination strategy to achieve specific stimulation of AS MNs by optogenetic tools: I) Silencing of optogenetic tool protein expression in the non-target subsets of MNs by dsRNA; II) Using the Q system for conditional expression. The transcriptional activator QF binds to the QUAS sequence to induce optogenetic tool expression. The transcriptional inhibitor QS suppresses expression in unwanted cells by binding to QF; III) Selective illumination of the VNC MNs by 470 nm blue light. The body of the worm was divided into 11 segments, of which 3–10 were illuminated in animals moving freely on agar plates. **B)** Expression pattern of ChR2(H134R)::YFP in AS MNs by the dsRNA subtractive approach; scale bar, 20 µm. **C)** Representative body postures kymograph (20 s) of normalized 2-point angles of a 100-point spine, calculated from head to tail of the animal. Positive and negative curvature is represented by blue and red color. Animal expressed ChR2 in AS MNs as in A I and was illuminated after 10 s as in A III. Blue bar, period of 470 nm illumination. **D)** Photodepolarization of AS MNs by ChR2 (in animals raised with ATR): I, II) Locomotion speed: Mean ± SEM crawling speed of animals before and during blue illumination (blue bar), comparing animals expressing ChR2 in AS MNs or in all types of cholinergic MNs in the VNC, raised in the presence or absence of ATR (II: Group data of mean speed of the animals before (15–20 s) and during (21–26 s) ChR2 photoactivation); III, IV) Mean ± SEM body length of the animals shown in I (IV: Group data of the mean length before (15–20 s) and during (21–26 s) photoactivation). **E)** Depolarization of subsets of AS MNs in body segments. I) Scheme of anterior, midbody and posterior segmental illumination; II) Representative body posture kymographs of 2-point angles from head to tail before (20 s) and during ChR2 photoactivation by blue light in the segments of the worm body, corresponding to experiments as in E I). See also Supplementary Video S1 and Supplementary Figure S1B. P values * ≤ 0.05; ** ≤ 0.01; *** ≤ 0.001. Number of animals is indicated in D. Statistical test in D II and IV: ANOVA with Tukey’s post hoc test.

### Depolarization of AS MNs activates body wall muscles (BWMs) and increases body bending during locomotion

*C. elegans* moves by propagating undulation waves along the body. Body bends are generated by cholinergic neurons mediating contraction of muscles on one side, and by GABAergic neurons mediating simultaneous relaxation of the contralateral side of the body (Donnelly et al., 2013; McIntire et al., 1993). According to the wiring diagram (Chen et al., 2006; Varshney et al., 2011; White et al., 1986) AS MNs send synapses mostly to the dorsal BWM cells (68, i.e. 47 % of all presynaptic contacts) and to inhibitory ventral (GABAergic) VD MNs (66 synapses, i.e. 46 %). To confirm that depolarization of the AS MNs evokes postsynaptic currents, we expressed and photo-activated channelrhodopsin-2 (ChR2(H134R); Nagel et al., 2005) in AS MNs, while recording electrically from dorsal muscles in dissected preparations (**Supplementary Fig. S1A**). We preserved commissural connections from the ventral nerve cord (where AS MN cell bodies reside), by cutting on the left side (i.e. opposite to where the AS commissures run). *C. elegans* MNs exhibit spontaneous activity leading to miniature post synaptic currents (mPSCs), but may generate rhythmic PSC bursts when triggered by PIN activation (Butler et al., 2014; Gao and Zhen, 2011; Kawano et al., 2011; Liewald et al., 2008; Liu et al., 2017; Schultheis et al., 2011; Wen et al., 2012). When activating AS MNs using ChR2, we observed a large peak current (ca. 400 pA), followed by mPSC firing at an increased rate for the duration of the illumination. This activity was similar to previously observed tonic activity after prolonged ChR2 activation of all cholinergic neurons (Liewald et al., 2008; Liu et al., 2009). Thus, depolarization did not induce any obvious intrinsic rhythmic activity in AS MNs.

Next, we measured parameters of crawling in intact worms moving freely on agar substrate. Activation of ChR2 in all cholinergic neurons including VNC MNs leads to strong contraction of the worm body and coiling (Zhang et al., 2007; Liewald 2008). AS MNs innervate only dorsal muscles, thus we wondered if their simultaneous depolarization would hinder propagation of the body bending wave. Animals in which ChR2 was activated in AS MNs kept the ability to propagate the undulation, yet they displayed a distorted wave, deeper bending (**Fig. 1C**), and transiently reduced speed (**Fig. 1DI, II**). Furthermore, photo-depolarization of AS MNs evoked body contraction, though this was reduced when compared to ChR2 activation of all VNC cholinergic MNs (**Fig. 1DIII, IV**). These behavioral phenotypes were blue-light dependent, and absent in transgenic animals raised without all-*trans* retinal (ATR), the obligate ChR2 co-factor. As the locomotion bending wave propagates from head to tail, we probed how AS MN activity contributes to this propagation. We thus stimulated AS MNs in small segments of the body (anterior, midbody, posterior; **Fig. 1EI**). These manipulations neither caused marked disruption of the wave (**Fig. 1EII**), nor did they reduce speed. However, they led to a reduction of body length (**Supplementary Fig. S1BI-VI**), most pronounced after stimulation of the anterior segment. In sum, AS MN depolarization facilitates, but may not play an instructive role in generating the undulatory wave.

### AS MN depolarization causes asymmetric BWM activation and a dorsal bias during locomotion

In contrast to the A and B class MNs, the AS MNs have no ‘opposing’ partner neurons (like VA/DA or VB/DB) and innervate (dorsal) BWMs and inhibitory VD neurons (that innervate ventral muscle). We wondered whether this evokes biased activation of dorsal BWMs, and thus used Ca^2+^ imaging (GCaMP3) in BMWs of immobilized animals (**Fig. 2AI**), while we photostimulated (ChR2) AS MNs. In animals raised with ATR, we observed asymmetric responses in the BWM during photoactivation of AS MNs: The Ca^2+^ signal in dorsal muscle cells increased, while it simultaneously decreased in the ventral muscles (**Fig. 2AII-IV, Supplementary Video S2**). In contrast, no such effect was observed in animals raised without ATR, i.e. containing non-functional ChR2: Here, when averaged over many animals, no net change in Ca^2+^ signals occurred, as spontaneously arising fluctuations canceled out. This functional asymmetry during AS MN photostimulation also affected locomotion, as these animals crawled in circles (**Fig. 2BI; Supplementary Video S1**). This was due to a bias towards the dorsal side, measured at the anterior body (**Fig. 2BIII-VI**), and leading to a mild, but significant increase in average bending along the body (**Fig. 2BV, VI**). In contrast, when all VNC cholinergic neurons were stimulated, animals showed deep bending with no dorsal or ventral bias, thus strongly slowing locomotion (**Figs. 1DI, II; 2BII, V, VI**). In sum, depolarization of AS MNs contributes to dorso-ventral coordination and likely facilitates navigation.

**Figure 2.**
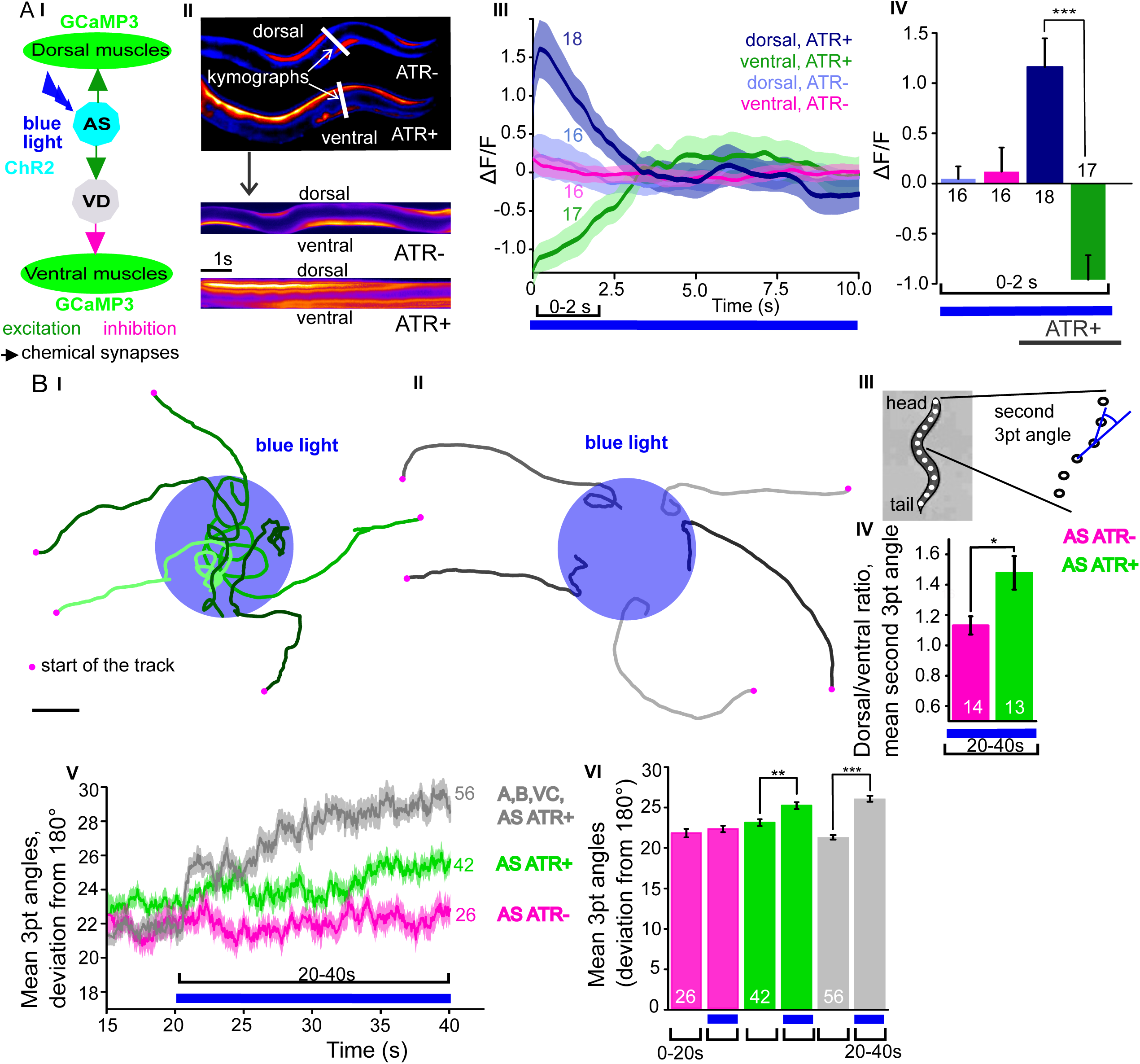
Photodepolarization of AS MNs causes activation of dorsal and simultaneous inhibition of ventral BWMs, and a dorsal bias in freely crawling animals. **A)** I) AS MNs expressing ChR2 are illuminated by 470 nm blue light, Ca^2+^ signal is recorded in the BWM expressing GCaMP3 (arrows, chemical synapses). II) Upper panel: Representative snapshots of Ca^2+^signals in BWM cells during blue light illumination in animals cultivated with and without all-trans-retinal (ATR); Lower panel: Kymographs of Ca^2+^ dynamics during 15 s of blue illumination, along the white lines indicated in the upper panel, covering both dorsal and ventral BWMs. III, IV) Mean Ca^2+^ signals (ΔF/F ± SEM) in dorsal and ventral BWM during the first 10 s of illumination (III) in animals raised with and without ATR and group data (IV), quantified during the first 2 s of illumination. B) I, II) Representative locomotion tracks of freely moving animals (raised with ATR) with ChR2 expressed only in AS MNs (I) or in all cholinergic MNs (II) before (20 s) and during photostimulation (20 s) by 470 nm blue light (indicated by blue shaded area; tracks are aligned such that they cross the blue area at the time of light onset). III) Schematic showing the thirteen points defining eleven 3-point angles along the spine of the animal. VI) Mean (± SEM) ratio of dorsal to ventral bending at the 2^nd^ 3-point bending angle in animals expressing AS::ChR2 during photostimulation (animals raised with and without ATR).V) Mean (± SEM) time traces of all 3-point bending angles before and during blue illumination (blue bar; ChR2 in AS MNs or in all cholinergic MNs; raised with and without ATR). VI) Group data as in V, comparing 20 s blue light illumination (blue bar), to the 20 s before illumination. See also Supplementary Video S2. P values * ≤ 0.05; ** ≤ 0.01; *** ≤ 0.001; number of animals is indicated. Statistical test in B III): T-Test; else: ANOVA with Tukey’s post-hoc test.

### AS MN ablation disrupts the locomotion pattern

The observed effects indicated an ability of AS MNs to evoke the bending wave during forward locomotion, and that they may play an important role in generating locomotion patterns. We probed the necessity of AS MNs for locomotion by ablation, as described earlier for other MNs and PINs (Chalfie et al., 1985; Gao et al., 2017; McIntire et al., 1993; Piggott et al., 2011). To this end, we used the genetically encoded, membrane targeted (*via* a pleckstrin homology -PH-domain) blue light activated <u>mini</u>ature <u>S</u>inglet <u>O</u>xygen <u>G</u>enerator (PH-miniSOG; Xu and Chisholm, 2016) and targeted illumination (**Fig. 3AI**). Brief illumination of the AS MNs with 470 nm light (2 mW/mm^2^, 2.5 min) led to visible and quantifiable locomotion defects: Animals with ablated AS MNs retained the ability to move, but crawled with lower speed, increased bending angles and an overall distorted undulation wave along the body, with a highly irregular pattern (**Fig. 3AII-IV, Supplementary Fig. S2A; Supplementary Video S3**). Thus, ablation of AS MNs disrupted coordination of the bending wave.

**Figure 3.**
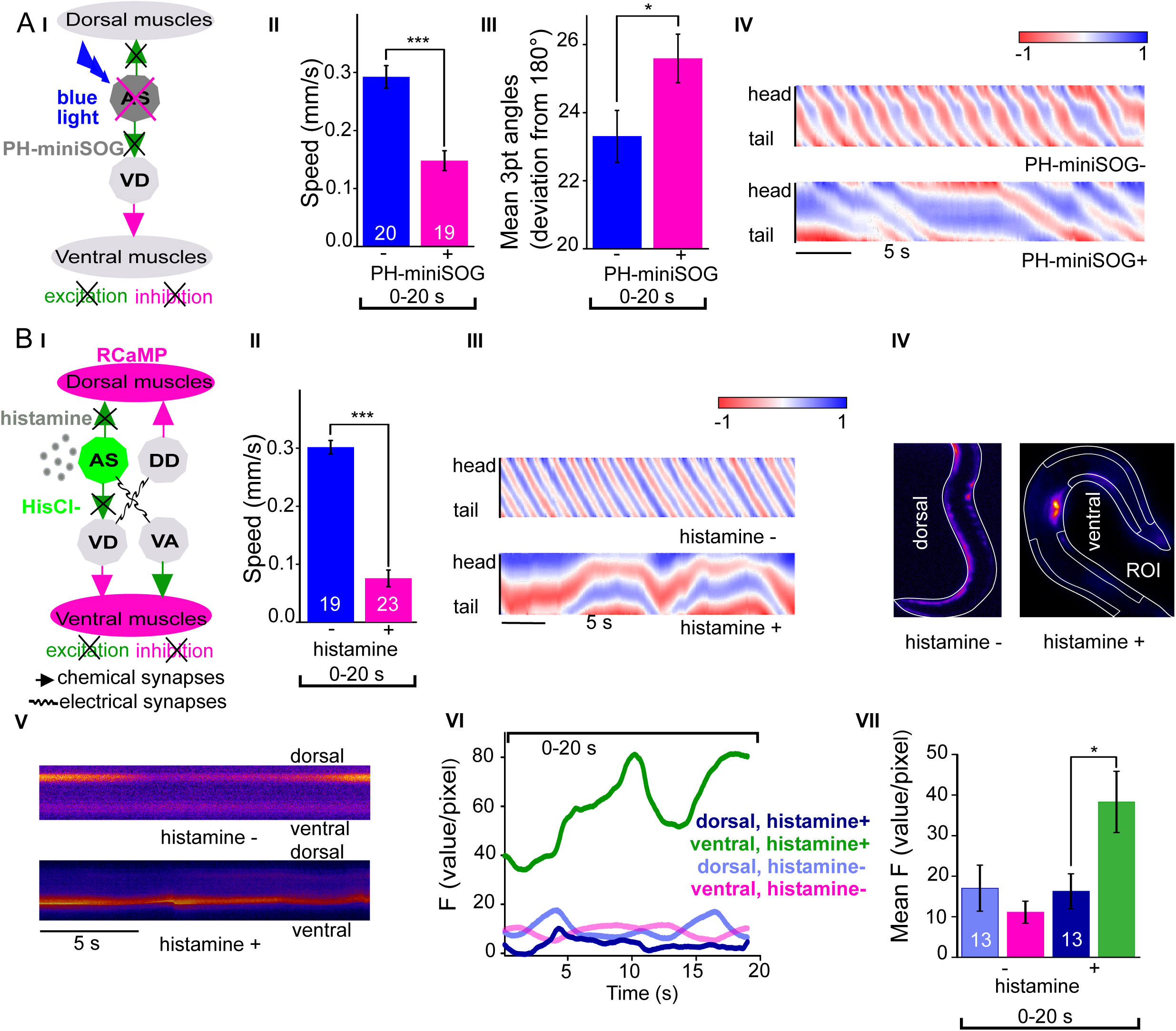
Optogenetic ablation and chronic hyperpolarization of AS MNs disrupts the locomotion pattern. **A)** I) Schematic of optogenetic ablation of AS MNs by PH-miniSOG and connectivity to relevant cell types (arrows, chemical synapses; curved lines, electrical synapses); Quantification of mean ± SEM speed (II) and bending angles (III) of animals without or with expression of PH-miniSOG (*via* the Q system) in AS MNs, following 150 s of blue light exposure and 2 h resting period; IV) Representative body posture kymographs (as in Fig. 1C) of wild type animal (upper panel) and animal expressing PH-miniSOG after photoactivation (lower panel). See also Supplementary Figure S3 and Supplementary Video 3. **B)** I) Schematic of Ca^2+^ imaging in BWM (RCaMP fluorescence) during hyperpolarization of AS MNs by HisCl1 (expressed in AS MNs *via* the Q system), and connectivity to relevant cell types (see also A I). II) Mean ± SEM speed of freely moving animals on agar dishes without and with 10 mM histamine. III) Representative body posture kymographs of animals freely moving on agar without (upper) or with 10 mM histamine (after 240 s incubation; lower panel). IV) Representative fluorescent micrographs of Ca^2+^ activity in the BWM of animals mounted on agar slides without (left) or with 10 mM histamine (after 240 s incubation; right panel). V) Representative kymographs (20 sec) of Ca^2+^ activity in dorsal and ventral BWM of animals as in IV. VI) Representative Ca^2+^ activity in dorsal and ventral BWM from animals as shown in IV, V. VII) Mean ± SEM fluorescence of dorsal and ventral BWM as in VI. See also Supplementary Fig. S2 and Supplementary video S4. P values* ≤ 0.05, *** ≤ 0.001; number of animals indicated in A II, III; B VII. Statistics: T-test for AII, III and B II; ANOVA with Tukey’s post-hoc test.for B VII.

### Chronic hyperpolarization of AS MNs eliminates Ca^2+^ activity in dorsal BWMs

Photodepolarization of AS MNs caused opposite effects on Ca^2+^ signals in dorsal and ventral muscles (**Fig. 2**). We wondered whether hyperpolarization of AS MNs may have reciprocal effects. AS MNs form excitatory chemical synapses to dorsal muscle and to VD MNs (the latter inhibit ventral muscle), but also gap-junctions (to VA MNs, exciting ventral muscle). Thus, several outcomes are conceivable: 1) Decrease of Ca^2+^ levels in dorsal, and increase in ventral muscles; 2) AS MN hyperpolarization may reduce ventral muscle activity via gap junctions to VA MNs; 3) A mixture of both, possibly causing oscillations. We thus used the *Drosophila* histamine-gated Cl^-^-channel HisCl1 (Pokala et al., 2014), as a hyperpolarizing tool (**Fig. 3BI**). Since *C. elegans* has no endogenous histamine receptors, HisCl1 can be specifically activated using histamine. First, we incubated animals expressing HisCl1 in AS MNs with histamine, and compared them to controls not incubated with histamine (**Fig. 3BII; Supplementary Fig. S2B; Video S4**). Animals on histamine plates moved significantly slower (ca. 75 % reduction) than animals without histamine, demonstrating that AS MNs are actively involved in promoting locomotion (however, this manipulation also affects other cholinergic neurons outside the VNC; see below). To analyze the possible reason for the reduced speed, we analyzed the crawling body postures (**Fig. 3BIII**). Histamine exposure strongly disturbed the propagation of the body wave, leading to very slow and irregular movement and frequent directional changes. To assess the effects of constant AS MN hyperpolarization on muscle physiology and activity, we expressed the red fluorescent Ca^2+^ indicator RCaMP in BWM cells (Akerboom et al., 2013), and analyzed spontaneous Ca^2+^ signals in ventral and dorsal muscles, in immobilized animals (**Fig. 3BIV**), either without or with histamine. Consistent with the dorsal innervation of muscles by AS MNs, spontaneous Ca^2+^ activity in animals with hyperpolarized AS MNs was observed only in ventral BWM, and animals showed ventral bending. Over time, on histamine, ventral Ca^2+^ fluctuations had much higher amplitude, while animals without histamine showed comparable and low amplitude fluctuations in both dorsal and ventral muscles (**Fig 3BV-VII**). In sum, hyperpolarization of AS MNs inhibits their excitatory signaling to dorsal muscles, and blocks their activation of GABAergic VD motor neurons, which in turn leads to ventral muscle disinhibition. This causes a strong bias to uniform ventral muscle activation, which likely disrupts propagation of the body wave.

### Acute hyperpolarization of AS MNs induces ventral muscle contraction through disinhibition

Chronic hyperpolarization of AS MNs by HisCl1 lacks temporal resolution, and, due to the expression from the *unc-17* promoter, despite our ‘subtractive’ expression, hyperpolarization of head and tail neurons could affect the outcome of these experiments. To avoid inhibition of these neurons, we looked for a potent hyperpolarizing optogenetic tool, enabling to use selective illumination for specific AS MN inhibition. We thus used the natural Cl^-^-conducting anion channel rhodopsin (ACR1), which causes strong (shunting) inhibition upon illumination (Sineshchekov et al., 2015; **Fig. 4A**).

**Figure 4.**
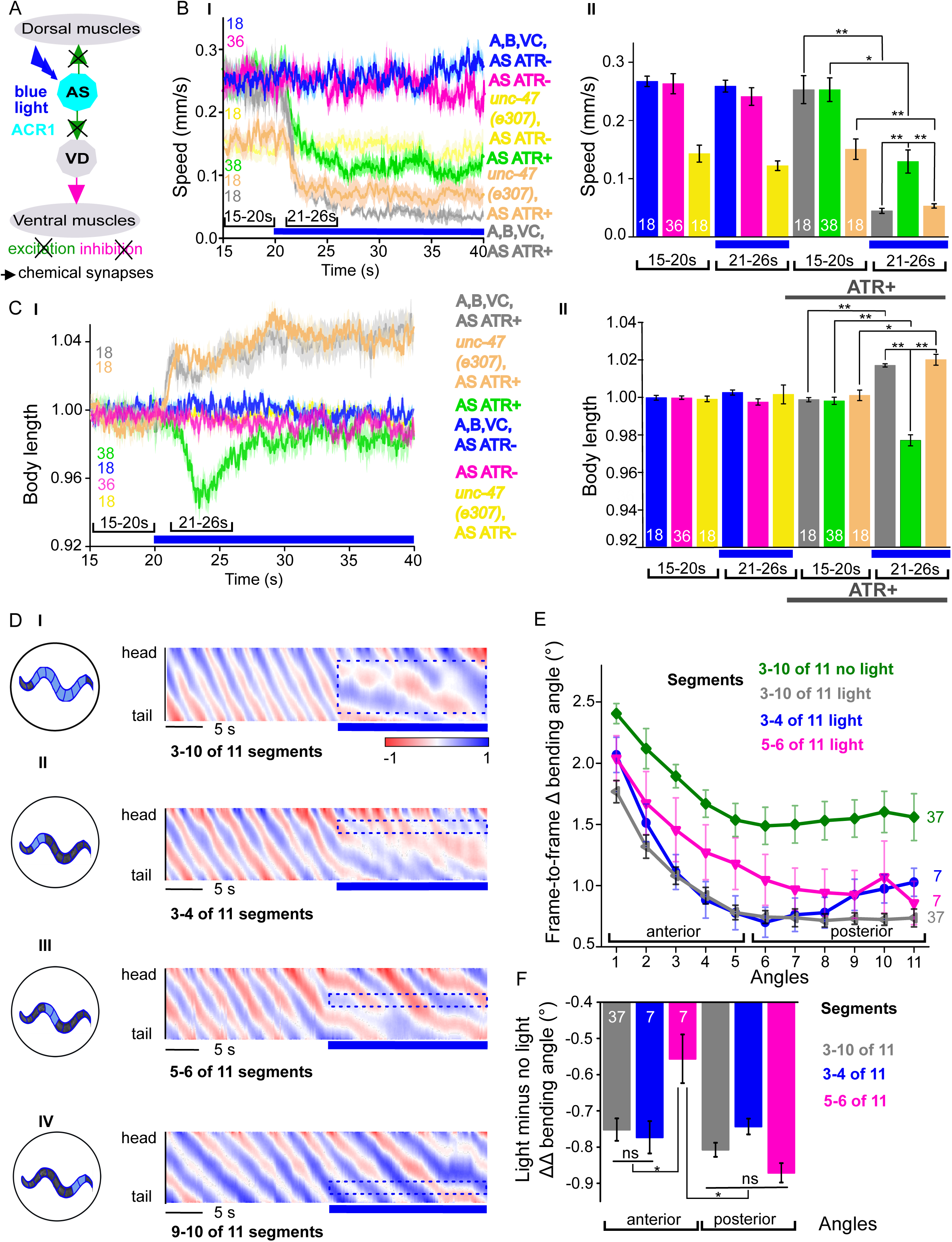
Acute optogenetic hyperpolarization of AS MNs ceases locomotion, causes disinhibition of ventral BWM via GABAergic VD MNs, and blocks propagation of the locomotion body wave: **A)** Schematic of experiment; hyperpolarization of AS MNs by the ACR1 anion channel rhodopsin activated by 470 nm blue light (arrows, chemical synapses). **B)** Time traces (I) and group data quantification (II) of mean ± SEM speed before (15–20 s) and during (21–26 s) blue illumination (indicated by blue bar). Compared are strains expressing ACR1 in all VNC cholinergic neurons, or in AS MNs only (*via* the Q system), in wild type or *unc-47(e307)* mutant background, raised in the presence or absence of ATR, as indicated. **C)** Time traces (I) and group data quantification (II) of mean ± SEM body length of the animals shown in B. **D)** Hyperpolarization of AS MNs in all (I), in the anterior (II), middle (III) and posterior (IV) segments of the worm body. Representative body postures kymographs of normalized 2-point angles from head to tail in animal expressing ACR1 in AS MNs before and during illumination by blue light in the indicated body segments. **E)** Mean, absolute difference of bending angles, from one video frame to the next (25 Hz), at each of eleven 3-point angles, for experiments as in (D). **F)** Mean difference of the differential bending angles between dark and illuminated conditions, for the analyses shown in (E). Data were averaged for the anterior 5 or the posterior 6 3-pt bending angles. See also Supplementary Fig. S3 and Supplementary Videos S5–8. P values * ≤ 0.05; ** ≤ 0.01; number of animals is indicated. Statistics: ANOVA with Tukey’s post hoc test.

Acute, ACR1-induced photo-hyperpolarization of all cholinergic neurons in freely moving animals strongly reduced crawling speed (to below 20 %) and essentially stopped locomotion (**Fig. 4B, C**). When we restricted expression of ACR1 and illumination to the AS MNs, we observed a similar reduction of speed, though not as pronounced (to ca. 35% of the initial speed; **Fig. 4B**). Controls (animals raised without ATR) showed no change in locomotion speed. These results, together with the HisCl1 experiments may suggest that the speed reduction was caused by a lack of ACh release from AS MNs to dorsal muscles. As this should cause partial relaxation of the body, we analyzed body length: For animals expressing ACR1 in all cholinergic neurons, we observed a prominent body elongation, in line with the absence of all excitatory (cholinergic) transmission to muscle (**Fig. 4C**). However, hyperpolarization of only the AS MNs led to partial and transient body contraction (**Fig. 4C, Supplementary Video S5**). This might be explained by synaptic connections of AS MNs to the GABAergic VD MNs: Hyperpolarization of AS MNs would reduce excitation of VD MNs, which in turn would cause dis-inhibition of muscle cells, and thus contraction. To test this, we repeated the experiment in *unc-47(e307)* mutants, lacking the vesicular GABA transporter, and thus GABAergic transmission. Consistently, *unc-47* mutants showed relaxation instead of contraction of BWMs (**Fig. 4C**). Body wave propagation was strongly attenuated, as for the analogous experiment using HisCl1 in AS MNs; however, using ACR1, this was induced within 2–3 s of illumination. Last, we probed if local AS MN inhibition (i.e. in anterior, midbody or posterior neurons) would block the propagation of the wave posterior from this point (**Fig. 4D**). This was the case: In about half of the animals tested, inhibition of anterior and midbody AS neurons hindered propagation of the wave to the posterior part of the body, leading to dragging behind of the tail region (**Fig. 4D, Supplementary Videos S6–8**). Analyses of the extent of movement in individual body segments showed that a reduction of movement was also found in the head region, however, this was more pronounced toward the posterior, particularly when the midbody AS neurons were inhibited (**Fig. 4E; Supplementary Fig. S3A** shows how the eleven 3-point angles analyzed correspond to illuminated body segments): The extent of reduction in body movement in the anterior part of the animal was significantly smaller than the change in posterior body movement (**Fig. 4F**). Animals also showed a reduction of speed, though not as pronounced as when all AS MNs were hyperpolarized, and length was not affected (**Supplementary Fig. S3BI-IV**). When the posterior segment was hyperpolarized, no obvious effects were observed. In sum, AS MNs are required for antero-posterior propagation of the body wave.

### Oscillatory AS MN activity correlates more strongly with body bends during forward crawling

Measuring Ca^2+^ transients in the ventral cord MNs during locomotion revealed higher activity states for B- and A-type MNs during forward and backward locomotion, respectively (Haspel et al., 2010; Kawano et al., 2011; Qi et al., 2013). Correlation of Ca^2+^ traces in AS MNs with dorsal body bends was previously shown in freely crawling animals (Faumont et al., 2011). Considering the unique situation of AS MNs, i.e. coupling with both forward and backward command interneurons, we wondered if AS MNs would maintain equal activity during both locomotion states. Thus, we measured Ca^2+^ transients in AS6 and AS7 in moving animals.

AS6 and AS7 showed oscillatory activity during locomotion, which was correlated with the change of body bends. During forward crawling, (anterior) AS6 activity preceded (posterior) AS7 activity by about 1–2 s (**Fig 5A, B; Supplementary Video S9**). To understand if AS MN Ca^2+^ transients are related to the locomotion body wave, we measured the angle defined by the position of AS6, the vulva, and AS7. We then performed cross-correlation analysis (for individual undulations, i.e. full periods of the bending wave) of the Ca^2+^ signal in AS6 or AS7 and the respective bending angle at the given time (**Fig. 5C, Supplementary Video S9**). Here, the Ca^2+^ signal in AS6 preceded the maximal bending at the vulva by about 2 s, while the signal in AS7 coincided (these correlations were moderate, but significant, the coefficients were ~0.35). Thus, the wave of activity in AS MNs appears to travel antero-posteriorly at the time scale of the undulatory wave (under a cover slip, slowing down locomotion; in animals moving freely on agar, the wave oscillates with ca. 0.5 Hz (Gao et al., 2017), while here, the delay of two maxima of undulation is ca. 3–4 s; **Fig. 5B**). We also measured cross-correlation between the Ca^2+^ signals in the AS6 and AS7 neurons (coefficient ~0.33). During reversal periods, as sometimes observed in our Ca^2+^ imaging experiments (**Fig. 5B**), there was essentially no correlation (coefficient ~ ± 0.1) of AS6 and AS7 Ca^2+^ signals with the vulva bending angles, or with each other, and there was also no obvious time lag between these signals (**Fig. 5D**). Yet, the peaks of AS6 and AS7 Ca^2+^ signals did not reveal any difference between forward and reverse movements, indicating that the cells were equally active during forward and reverse locomotion (**Fig 5E**). In sum, AS MNs showed oscillatory activity that was more strongly correlated with body bends during forward than during backward crawling.

**Figure 5.**
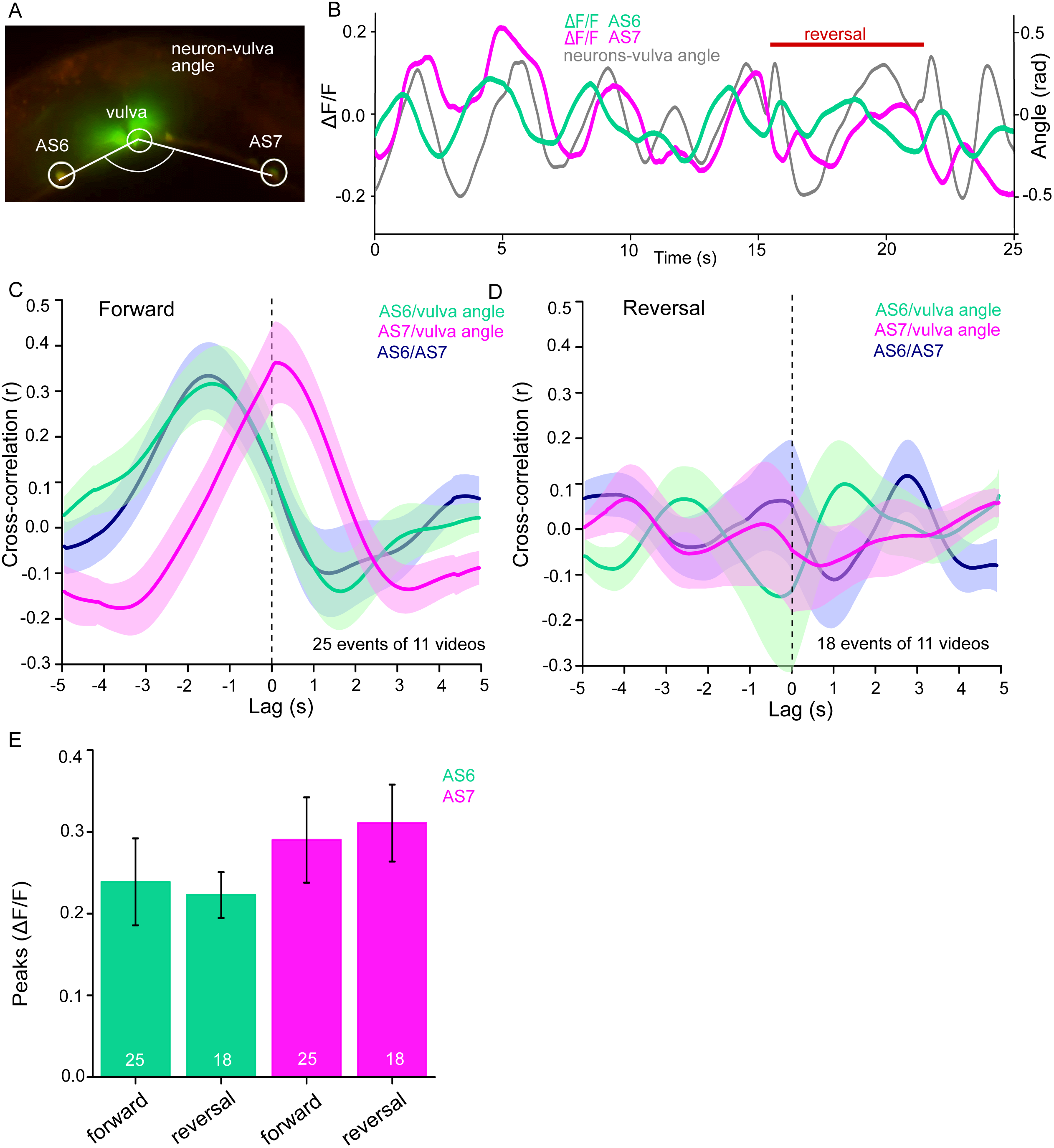
AS MNs show oscillatory Ca^2+^ activity in moving animals: **A)** Fluorescent micrograph (red and green as merged fluorescence channels) of the vulva region, showing red (mCherry) and green (GCaMP6), expressed in AS MNs (with the use of Q system), and GFP, expressed in vulva muscles. Angle between vulva and the two flanking AS6 and AS7 neurons is indicated. **B)** Representative analysis of time traces (25 s) of Ca^2+^ signals (ΔF/F) in AS6 and AS7, as well as the angle defined by the vulva and the two neurons during forward crawling, with a single reversal event (red bar). **C, D)** Cross-correlation analysis (mean ± SEM) of single periods of the body wave (5 s each) for each of the AS6 and AS7 GCaMP6 signals with the vulva angle, as well as for the two Ca^2+^ signals, during forward (C) or backward (D) locomotion. **E)** Comparison of the peaks Ca^2+^ signals (mean ± SEM) in AS6 and AS7, during forward or reverse locomotion, respectively. See also Supplementary Video S9. Number of animals is indicated in C-E. Statistical test: ANOVA with Tukey’s post-hoc test.

### AS MNs integrate signaling from both forward and backward premotor-interneurons

The PINs AVA, AVD, and AVE connect to the DA and VA MNs, and induce reversals and backward locomotion. Conversely, the PINs AVB and PVC are connected to the DB and VB forward MNs and mediate forward locomotion (Chalfie et al., 1985; Chronis et al., 2007; Kawano et al., 2011; Piggott et al., 2011; Qi et al., 2013; Wicks et al., 1996). Endogenous as well as stimulated activity of the PINs modulates activity of A- and B-type MNs (Kawano et al., 2011; Liu et al., 2017).

The AS MNs are postsynaptic for both backward (synapse number: AVA - 63, AVE - 7) and forward PINs (AVB - 13, PVC - 2). This suggests a bias of AS MNs for backward locomotion; however, as synapse number is not the only determinant of synaptic weight, also the opposite (as indicated by our AS MN Ca^2+^ imaging data) is conceivable. No chemical synapses are known from AS MNs towards the PINs, yet, there are 37 gap junctions reported between AVA and the AS MNs as well as 5 gap junctions between AVB and the AS MNs. Electrical synapses could mediate anterograde as well as retrograde signaling between AS MNs and PINs (Chen et al., 2006; Varshney et al., 2011; White et al., 1986). To assess whether depolarization of AVA and AVB would lead to observable and/or different Ca^2+^ responses in the AS MNs, we generated strains expressing ChR2 in the PINs and GCaMP6 in AS MNs (**Fig. 6AI**): One strain specifically expressed ChR2 in AVA (Schmitt et al., 2012) and another strain expressed ChR2 from the *sra-11* promoter in AIA, AIY, and AVB neurons, of which only AVB has direct synaptic connections to AS MNs. The respective animals were photostimulated and Ca^2+^ transients were measured in AS3 (anterior) and AS8 (posterior) MNs of immobilized animals, raised either in absence or presence of ATR (i.e. without and with functional ChR2). Stimulation of AVA or AVB both resulted in a steady, synchronous increase of the Ca^2+^ signal in the AS3 neuron; however, no increase was observed in animals raised without ATR (**Fig. 6AII-IV; Supplementary Videos S10, S11**). A similar increase was found in the AS8 neuron, and both AS3 and AS8 showed a synchronized increase of activity (**Supplementary Fig. S4A, B**). Thus, signaling from both forward and backward PINs is excitatory to the AS MNs. However, there was a clear difference in the response of the AS MNs to AVA vs. AVB stimulation, as depolarization of AVA produced a response of comparably low amplitude (up to 20 % ΔF/F after 3 s), while depolarization of AVB caused a response of significantly higher amplitude (up to 40 % ΔF/F; **Fig. 6AIV**). This could be due to differences in synaptic strength and/or ratio of chemical and electrical synapses between AVA-AS and AVB-AS, and may represent a physiological correlate of the apparently different involvement of AS MNs in forward vs. backward locomotion (**Fig. 5C, D**). Such inequality in regulated behavior based on imbalances in wiring was also observed for the PINs and A- and B-class MNs (Kawano et al., 2011; Roberts et al., 2016).Thus, despite predominant chemical synaptic connections between AVA and AS, AVB depolarization had stronger effects on AS MNs.

**Figure 6.**
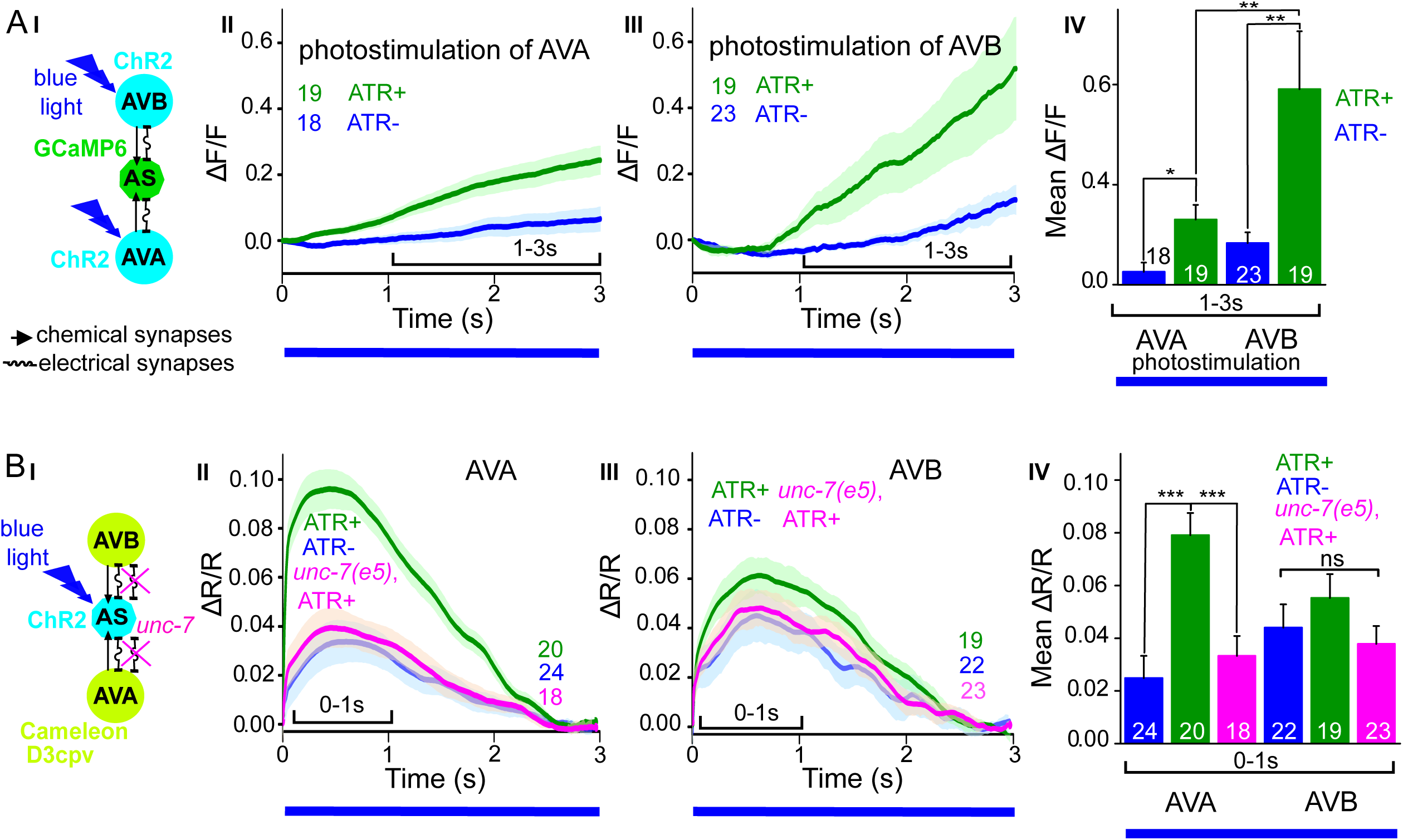
Reciprocal and asymmetric mutual activation of AS MNs and forward and reverse PINs, AVB and AVA. **A)** I) Schematic of the experiment for measurement of AS MN Ca^2+^signals (GCaMP6) during AVB or AVA photodepolarization *via* ChR2 with 470 nm blue illumination (arrows, chemical synapses; curved lines, electrical synapses). II, III) Time traces of mean (± SEM) Ca^2+^ transients (ΔF/F) in AS MNs during depolarization of AVA (II) and AVB (III) by ChR2, in animals raised in absence or presence of ATR. Brackets indicate time periods used for statistical analysis in IV. IV) Group data quantification of data shown in II and III (for the 1–3 s time period). See also Supplementary Fig. S5 and Supplementary Videos S10, 11. **B)** I) Schematic of the experiment for measurement of Ca^2+^ signals (cameleon) in AVB or AVA PINs during AS MN photodepolarization *via* ChR2 with 470 nm blue illumination. II, III) Mean (± SEM) of Ca^2+^ transients (ΔR/R YFP/CFP ratios) in AVA (II) and AVB (III) during AS MN depolarization, in wild type or *unc-7(e5)* mutant animals, raised in absence or presence of ATR. IV) Group data quantification of experiments in II and III (for the 0–1 s time period). P values * ≤ 0.05; ** ≤ 0.01; *** ≤ 0.001; number of animals is indicated. Statistical test: Mann-Whitney U test.

### Retrograde electrical signaling from AS MNs depolarizes AVA but not AVB interneurons

Recent observations revealed that AVA coupling to A-type MNs via gap junctions is strongly rectifying towards AVA (Liu et al., 2017). We thus wanted to explore the role of the electrical synapses between the PINs and the AS MNs in more detail, e.g. whether AS MN photostimulation could lead to depolarization of the PINs.

We generated strains expressing the ratiometric Ca^2+^ indicator cameleon (bearing CFP and YFP moieties; Miyawaki et al., 1997) in the command interneurons (driven by *sra-11* and *nmr-1* promoters, for expression in AVB and AVA, respectively) together with ChR2 expressed in the AS MNs (**Fig. 6BI**). Both promoters express in several head neurons, yet we could identify AVB and AVA by their position with respect to anatomical landmarks and with respect to other (known) fluorescent neurons. Photodepolarization of the AS MNs (in animals raised in the presence of ATR) caused Ca^2+^ transients in AVA (ΔR/R ~10%), but had no significant effect on Ca^2+^activity in AVB interneurons (**Fig. 6BII-IV**). A small (ΔR/R ~3–4%), insignificant increase of the ratio of the CFP/YFP signal was observed in the control animals raised without ATR, in both AVA and AVB. AS MNs express UNC-7 and INX-3 as the sole innexins. INX-3 is widely expressed in multiple tissues, and AVA and AVB also express UNC-7 (Altun et al., 2009; Starich et al., 2009). Thus, we used an *unc-7(e5)* null mutant, in which no electrical coupling should occur between AS MNs and AVA or AVB, and repeated the above experiments: The Ca^2+^ signal (ΔR/R ~4%) was now comparable to the signal observed in the control without ATR, indicating that UNC-7 electrical synapses are responsible for transmission between AS MNs and AVA. For AVB, we did not observe any significant effect in the *unc-7(e5)* mutant.

## DISCUSSION

Movement by undulations is remarkably effective across scales and in a variety of environments (Cohen and Sanders, 2014). Despite the diversity of their anatomy, the nervous systems of distantly related organisms may adopt similar strategies to control locomotion by undulations. Based on physiological data we revealed several features of the AS MNs highlighting their function in one of the most studied locomotion circuits, the VNC of *C. elegans*. The main findings of this work are: 1) Depolarization of AS MNs does not disrupt locomotion, but causes a dorsal bias. 2) AS MN hyperpolarization inhibits locomotion and prevents generation and propagation of the undulatory wave. 3) AS MN activity during locomotion is oscillatory, and is correlated more with forward than with backward locomotion. 4) AS MNs are stimulated by premotor interneurons, where the forward PIN AVB exerts stronger activity in AS MNs than the backward PIN AVA. 5) AS MNs exhibit functional electrical connections to the backward PIN AVA. Our findings for AS MNs in the *C. elegans* locomotor circuit have parallels in several animal models (see below).

### AS MNs act in coordination of dorso-ventral bends, antero-posterior wave propagation and possibly forward-backward states

AS MNs occupy a significant part of the VNC circuit (11 of 75 neurons) and two AS MNs are present in each functional segment of the circuit (Haspel and O’Donovan, 2011). Yet, in absence of physiological information, they were missing in many models representing the locomotor circuit function in *C. elegans* (Von Stetina et al., 2005; Zhen and Samuel, 2015). We showed that AS MNs are important for several aspects of locomotion, among them dorso-ventral coordination: Their depolarization induces postsynaptic currents in dorsal BWMs leading to contraction and dorsal bias in freely moving animals, while AS MN hyperpolarization eliminates activity in dorsal BWMs and induces contraction of contralateral ventral BWMs through disinhibition. Thus, it is likely that AS MNs counteract neurons providing a ventral bias, e.g. the VA and VB MNs, or even the VC neurons (Faumont et al., 2011; White et al., 1986). This corresponds to recent computational studies, which predicted a significant role of AS MNs in coordinating dorso-ventral bending (Olivares et al., 2017) and in the control of BWMs (Yan et al., 2017). Furthermore, AS MNs are active both during forward and reverse locomotion, however, correlation of their activity with movement is strong only for forward locomotion. In line with this, AS MN inhibition disrupts propagation of the antero-posterior body wave. Since the AS MNs connect to both forward and backward PINs, they could play a role in integrating forward and backward locomotion motifs, e.g. by providing an electrical sink (or source) for the PINs of the respective opposite direction (**Fig. 7A, B**). Similar functions were shown for A-type MNs and AVA (Kawano et al., 2011; Liu et al., 2017) as well as for V2a interneurons and MNs in zebrafish (Song et al., 2016).

**Figure 7.**
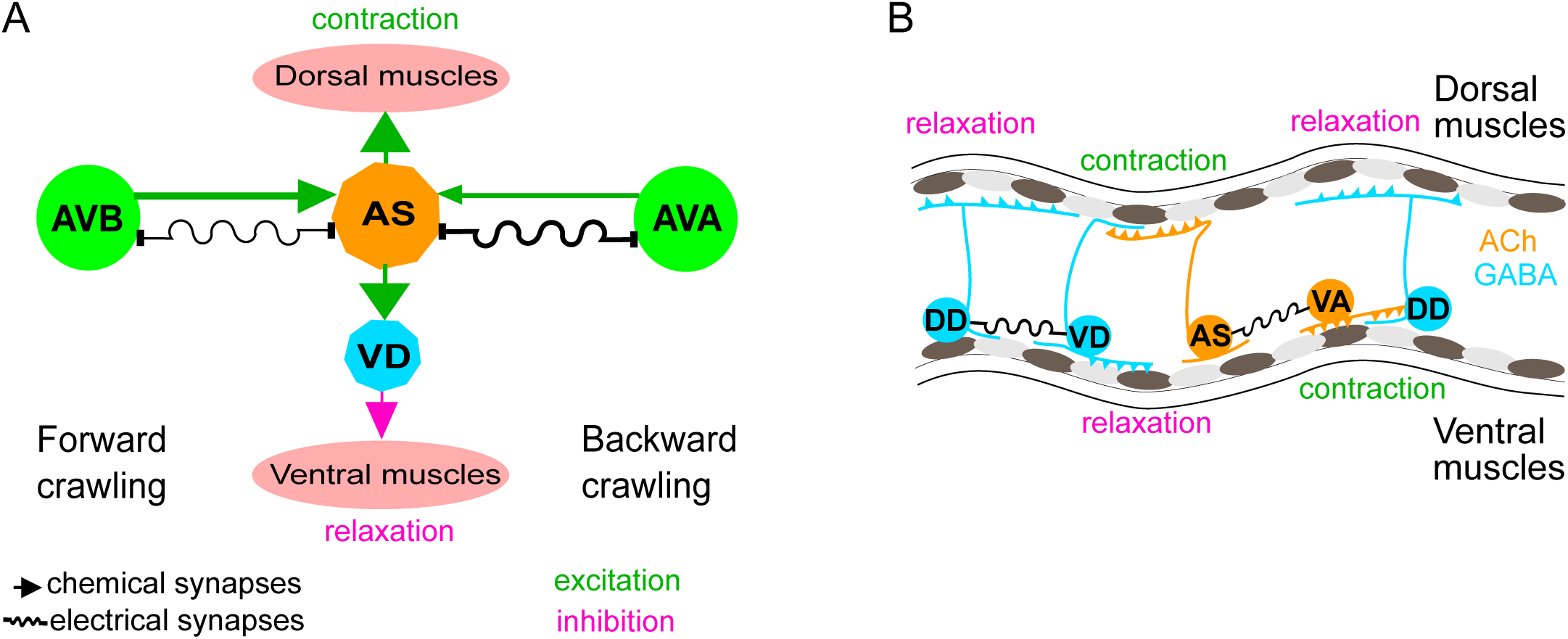
Models summarizing findings of this work: **A)** AS MNs control dorso-ventral bending coordination during forward and backward locomotion, by excitatory chemical transmission to dorsal muscles and ventral GABAergic VD MNs, thus causing ventral inhibition. Interconnections (arrows, chemical synapses; curved lines, electrical synapses; thickness of lines indicates relative synaptic strength) of AS MNs and their other synaptic partners, i.e. the PINs AVB and AVA, via both chemical synapses from the PINs and (reciprocal) electrical synapses from AS MNs are also shown. Data in this work suggest strong (chemical) excitatory regulation of AS MNs by AVB during forward locomotion, and reciprocal electrical regulation of AVA by AS MNs. **B)** Interconnections and functional roles of AS MNs and other VNC MNs during the propagation of the undulatory wave along the body. Depolarization (which could be initiated by AVB, not shown here, or by proprioceptive feedback from the adjacent body segment) of AS MNs causes contraction of the dorsal BWMs and simultaneous relaxation of ventral BWMs through the excitation of VD MNs. This phase is followed by contraction of ventral BWMs, e.g. through the electric coupling of AS and VA MNs, and relaxation of the dorsal BWMs through VD-DD electrical coupling or VA-DD chemical synapses. Cholinergic (orange) and GABAergic (blue) cell types are indicated. Antero-posterior localization of cell bodies and connectivity to other cell types are arbitrary.

AS MNs as other MN types innervate only one side of the BWMs (dorsal). However, unlike other MN classes, AS MNs have no obvious class of ‘partner’ neurons innervating only ventrally (of the VC neurons, which innervate ventral muscle, there are only six, and two of them mainly innervate vulval muscle). AS MNs thus provide asymmetric excitation, which may be required to enable complex regulatory tasks like changing direction during navigation. Indeed, optogenetic depolarization of AS MNs, resulting in curved locomotion tracks, mimicked the ‘weathervane’ mode of navigation towards a source of attractive salt (Iino and Yoshida, 2009). During locomotion, higher order neurons, that integrate sensory information, might influence the AS MNs to generate this bias to the dorsal side. In the lamprey, lateral bends were shown to be caused by asymmetry in stimulation of the mesencephalic locomotor region (Sirota et al., 2000), and in the freely moving lamprey even comparatively small left or right asymmetries in activity of the reticulospinal system corresponded to lateral turning (Deliagina et al., 2000).

Asymmetry in contralateral motifs of complex locomotor circuits is also known from vertebrate spinal cord CPGs in flexor-extensor coordination (Grillner and Wallén, 2002). In mice, flexor motor neurons are predominantly active and inhibit extensor motor neurons, which in turn show intervals of tonic activity between inhibitory states, corresponding to flexor bursts (Machado et al., 2015; Rybak et al., 2015). When comparing numbers of synaptic inputs from *C. elegans* cholinergic MNs to BWM (Varshney et al., 2011; White et al., 1986), predominance is apparent in excitatory neuromuscular junctions from A- and B-type MNs to ventral muscles, as well as in the corresponding contralateral synapses to inhibitory DD MNs, which innervate dorsal muscle. Therefore, tonic activity of B- or A-type MNs would be expected to generate a bias towards ventral bending, and this could be balanced by excitation of AS MNs. In addition, VC neurons may contribute in counteracting AS MN function (see above). However, the compressed nature of the *C. elegans* nervous system, in which single neurons fulfill multiple tasks that in higher animals are executed by layers of different cells, may not always allow for the direct comparison to vertebrate systems.

The groups of forward (AVB, PVC) and backward PINs (AVA, AVD, AVE), respectively, are synchronized (Prevedel et al., 2014; Schrödel et al., 2013), inhibit each other and change their state stochastically (Pierce-Shimomura et al., 2008; Roberts et al., 2016). Despite anatomical prevalence of chemical synapses from AVA to AS MNs, our data showed larger AS MN effects after stimulation of the forward PIN AVB, and a stronger correlation of AS MN activity with forward locomotion. Recently, ability of MNs to modulate activity of PINs was shown in several animal models: for B-type MNs changing the inhibitory chemical transmission of AVB to AVA in *C. elegans* (Kawano et al., 2011; Liu et al., 2017); for MNs regulating the frequency of crawling in *Drosophila* (Matsunaga et al., 2017); for MNs affecting activity of the excitatory V2a neurons in zebrafish (Song et al., 2016); and, in the mouse, such activity was suggested for MNs, changing the frequency of rhythmic CPG activity after stimulation by rhodopsins (Falgairolle et al., 2017). Positive feedback from MNs required the function of gap junctions, coupling between MNs and PINs in all these systems. Our data suggest that AS MNs are more relevant during forward locomotion, and receive stronger inputs from the forward PIN AVB, while their electrical feedback is stronger to the backward PIN AVA. If the latter would provide inhibition in the context of the free moving animal, e.g. as an electrical sink for AVA, AS MN activity should exert a bias to promote the forward locomotion state.

### AS MNs as CPGs?

CPGs are dedicated neural circuits with intrinsic rhythmic activities (Grillner, 2006; Guertin, 2013). In many organisms including those showing undulatory movement (e.g. leech, lamprey), series of CPGs are distributed along the length of the body in locomotor neural circuits (Kristan et al., 2005; Mullins et al., 2011). In *C. elegans*, the bending wave can be generated even in the absence of all PINs (Gao et al., 2017; Kawano et al., 2011; Zheng et al., 1999) as well as in absence of GABAergic MNs (Donnelly et al., 2013; McIntire et al., 1993). The existence of series of CPGs in the *C. elegans* VNC was discussed for a long time, and single neurons or small groups of neurons were suggested (Cohen and Sanders, 2014; Zhen and Samuel, 2015). B-type MNs are able to propagate the bending wave posteriorly from a likely head CPG that generates head oscillations (Hendricks et al., 2012; Shen et al., 2016), simply by proprioceptive coupling (Wen et. al., 2012). Recently, pacemaker properties of the posterior A-type MN, DA9, were revealed during backward locomotion, that are based on the activity of a P/Q/N-type Ca^2+^ channel (Gao et al., 2017). Computational modelling of repeating units of the VNC, based on connectome data, identified a dorsally oriented sub-circuit consisting of AS, DA, and DB MNs, which could act as a potential CPG (Olivares et al., 2017).

### Potential gating properties of AS MNs

Among PINs, AVB and AVA are most important for enabling forward and backward locomotion, respectively (Chalfie et al., 1985; Kato et al., 2015; Roberts et al., 2016). Bistable states with two distinct membrane potentials, i.e. up and down states, that ‘gate’ activity of the downstream target neurons, were shown for several interneurons including AVA and AVB (Gordus et al., 2015; Kato et al., 2015; Mellem et al., 2008). For MNs, bistability was inferred for the A-, B- and D- types of MNs from direct recordings (Liu et al., 2014) as well as from the graded responses in muscles, corresponding to shorter or longer activity bursts in MNs (Liu et al., 2017). Further, all-or-none responses in BWM cells, corresponding to spiking neurons as well as to mammalian skeletal muscles, that result from integrating graded excitatory and inhibitory input from MNs, were demonstrated (Gao and Zhen, 2011; Liu et al., 2011). AS MNs may similarly integrate inputs from forward and backward PINs, or themselves influence the PINs via UNC-7 and/or INX-3 gap junctions, to gate signal propagation in the VNC during forward locomotion, or to couple to A-type MN oscillators (DA9), via the AVA PIN during backward locomotion (Gao et al., 2017). In line with this hypothesis, we found ceasing of locomotion when AS MNs were hyperpolarized. The different AS MN responses to depolarization of AVB and AVA could represent biased activation, analogous to the imbalanced activities of A- and B-type MNs during forward and reverse locomotion (Kawano et al., 2011). Gating neurons that affect rhythmic properties of CPGs are also known for the leech locomotor circuit (Friesen and Kristan, 2007; Mullins et al., 2011; Taylor et al., 2000).

### Conclusions

The previously uncharacterized class of AS motor neurons is specialized in coordination of dorso-ventral undulation bends during wave propagation, a feature maintained by asymmetry in both synaptic input and output. Moreover AS neurons integrate signals for forward and reverse locomotion from premotor interneurons and potentially gate ventral nerve cord CPGs via gap junctions.

## EXPERIMENTAL PROCEDURES

### Strains and Genetics

*C. elegans* strains were maintained under standard conditions on nematode growth medium (NGM) and fed by *E. coli* strain OP50–1 (Brenner, 1974). Transgenic lines were generated using standard procedures (Fire and Pelham, 1986) by injecting young adult hermaphrodites with the (plasmid-encoded) transgene of interest and a marker plasmid that expresses a fluorescent protein. In some cases, empty vector was included to increase the overall DNA concentration to 150–200 ng/µl.

The following strains were used or generated for this study: **N2** (wild type isolate, Bristol strain), **CB5**: *unc-7(e5)X*, **CB307**: *unc-47(e307)III*, **CZ16469:** *acr-2(n2420)X; juEx4768[psra-11::ChR2::yfp]* (Qi et al., 2013), **PD4665**: *wt; ccIs4655[pes-10::GFP;dpy-20+]*, **RM2558**: *wt; Is[punc-17::GFP-NLS]*, **ZM5091**: *wt; hpIs190[pnmr-1(short2)-D3cpv; lin-15+]*, **ZM5089**: *unc-7(e5)X*; *hpIs190*, **ZM5132**: *wt; hpIs179[psra-11-D3cpv]*, **ZM5136**: *unc-7(e5)X*; *hpIs179* (all ZM strains are kind gift from Mei Zhen), **ZX460**: *wt; zxIs6[punc-17::ChR2(H134R)::yfp; lin-15+]V*, **ZX499**: *wt; zxIs5[punc-17::ChR2(H134R)::yfp; lin-15+]X*, **ZX1023**: *lite-1(ce314)X; zxIs30[pflp-18::flox::ChR2mCherry::SL2::GFP; pgpa-14::nCre; lin-15+]*, **ZX1396**: *wt; zxIs51[pmyo-3::RCaMP1h]*, **ZX2002**: *lite-1(ce314)X; zxIs6*, **ZX2004**: *lite-1(ce314)X; zxEx1016 [punc-4::ChR2_RNAi_sense & antisense_ pmyo-2::mCherry]; zxEx1017[pacr-5::ChR2_RNAi_sense & antisense_; pmyo-3::mCherry]*, **ZX2007**: *wt; zxIs5*; *zxEx1016; zxEx1017*, **ZX2008**: *wt; zxEx1023[punc-17::QF; pacr-5::QS::mCherry; punc-4::QS::mCherry; QUAS::ACR1::YFP; pmyo-2::mCherry]*, **ZX2011**: *wt; zxEx1020[punc-17::QF; pacr-5::QS::mCherry; punc-4::QS::mCherry; QUAS::HisCl1::GFP; pmyo-2::mCherry]*, **ZX2012**: *lite-1(ce314)X; ccIs4655[pes-10::GFP; dpy-20+]; zxEx1021[punc-17::QF; pacr-5::QS::mCherry; punc-4::QS::mCherry; QUAS::GCaMP6::SL2::mCherry; pmyo-2::mCherry]*, **ZX2110**: *wt; mdIs[punc-17::GFP-NLS;] zxEx1024[punc-17::QF; pacr-5::QS::mCherry; punc-4::QS::mCherry; QUAS::PH-miniSOG; pmyo-2::mCherry]*, **ZX2113**: *unc-47(e307)III; zxEx1029[punc-17::QF; pacr-5::QS::mCherry; punc-4::QS::mCherry; QUAS::ACR1::YFP; pmyo-2::mCherry]*, **ZX2114**: *wt; zxIs51; zxEx1020*, **ZX2127**: *lite-1(ce314)X; zxIs30; zxEx1021*, **ZX2128**: *lite-1(ce314)X; juEx4768; zxEx1021*, **ZX2132**: *wt; zxIs5*; *zxEx1016, zxEx1017; zxEx1028[pmyo-3::GCaMP3]*, **ZX2212**: *lite-1(ce314)X; hpIs179; zxIs6; zxEx1016, zxEx1017*, **ZX2213**: *lite-1(ce314)X; hpIs190; zxIs6; zxEx1016, zxEx1017*, **ZX2217***: unc-7(e5)X; hpIs190; zxIs6; zxEx1016, zxEx1017;* **ZX2220**: *unc-7(e5); hpIs179; zxIs6; zxEx1016, zxEx1017*, **ZX2221**: *unc-7(e5)X; zxIs6; zxEx1016, zxEx1017*

### Molecular biology

We used the following promoters: 3.5-kb *punc-17* (Sieburth et al., 2005), 2.5 kb *punc-4* (Miller et al., 1992) and 4.3 kb *pacr-5* (Winnier et al., 1999), genomic DNA sequence upstream of the ATG start codon of each gene, respectively. The *punc-4*::ChR2 sense and antisense construct was generated as follows: The p*unc-4* promoter from plasmid pCS139 was subcloned into pCS57 (Schultheis et al., 2011) by *Sph*I and *Nhe*I, 1 kb antisense sequence was amplified from the ligated construct pOT1 *punc-4*::ChR2::YFP using primers (5’-GGGGTTTAAACAGCTAGCGTCGATCCATGG-3’ and 5’-CCCGCGGCCGCCCAGCGTCTCGACCTCAATC-3’) and subcloned into the same construct by *Not*I and *Pme*I to get pOT2. To silence ChR2 expression under the *pacr-5* promoter we used a sense and antisense strands approach (Esposito et al., 2007) as follows: The *pacr-5* promoter was amplified from genomic DNA using primers (5’-TTATGATGCGAAAGCTGAATCGAGAAAGAG-3’, 5’-CCATGCTTACTGCACTTGCTTCCCATACTTC-3’, nested 5’-GGGGCATGCATCGAGAAAGAGAAGCGGCG-3’, 5’-CCCGCTAGCAAAGCATTGAAACTGGTGAC-3’) and subcloned into pCS57 with *Sph*I and *Nhe*I to yield pOT3 (*pacr-5*::ChR2::YFP). The sense and antisense strands were amplified from this construct using primers (for the coding region of ChR2: 5’-ATGGATTATGGAGGCGCCC-3’, 5’-CCAGCGTCTCGACCTCAATC-3’; for the promoter sense: 5’-GGCGGAGAGTAGTGTGTAGTG-3’ and 5’-GGGCGCCTCCATAATCCATCAAAGCATTGAAACTGGTGACGAG-3’; for the promoter antisense: 5’-GGCGGAGAGTAGTGTGTAGTG-3’ and 5’-GATTGAGGTCGAGACGCTGGCAAAGCATTGAAACTGGTGACGAG-3’; for fusion of sense strand: 5’-GCGGTTTCACGCTCTGATGAT-3’ and 5’-CTCAGTGCCACCAATGTTCAA-3’; and for fusion of the antisense strand: 5’-GCGGTTTCACGCTCTGATGAT-3’ and 5’-GCGCGAGCTGCTATTTGTAA-3’).

The pOT6 *punc-17*::QF construct was generated as follows: QF::SL2::mCherry sequence was amplified from plasmid XW08 (a kind gift from Kang Shen and Xing Wei) using primers 5’-CAGGAGGACCCTTGGATGCCGCCTAAACGCAAGAC-3’ and 5’-AGTAGAACTCAGTTTTCTGATGACAGCGGCCGATG-3’, and subcloned into pCS57 (Schultheis et al., 2011) using In-Fusion cloning (Takara/Clontech). The SL2::mCherry fragment was cut out by *Sal*I and *Bbv*CI and 5’-overhangs were filled in with Klenow polymerase (NEB). The pOT8 *pacr-5*::QS::mCherry construct was generated by subcloning the *pacr-5* promoter from pOT3 into vector XW09 (Wei et al., 2012) (a gift from Kang Shen and Xing Wei) with *Sph*I and *NheI*. The pOT7 *punc-4*::QS::mCherry construct was generated by subcloning the *punc-4* promoter from the pOT1 plasmid into XW09 vector with *Sph*I and *NheI*. The pOT10 pQUAS::Δpes-10::HisCl1::GFP construct was generated by subcloning the sequence encoding HisCl1::GFP from plasmid pNP403 (Pokala et al., 2014) (kind gift from Navin Pokala and Cori Bargmann) into the XW12 vector (a gift from Kang Shen and Xing Wei) with *Asc*I and *Psp*OMI. The pOT11 pQUAS:: Δpes-10::GCamP6::SL2::mCherry construct was generated as follows: pQUAS::Δpes-10 sequence was amplified from plasmid XW12 using primers 5’-ACAGCTATGACCATGATTACGCCAAG-3’ and 5’-CCCCGCGGCCGCCCAATCCCGGGGATCCTCTA-3’, and subcloned into plasmid p*lin-11*::GCaMP6::SL2::mCherry with *Sph*I and *Not*I. The pOT13 pQUAS:: Δpes-10::ACR1::YFP construct was generated as follows: ACR1::YFP sequence was amplified from plasmid pAB03 (*punc-17*::ACR1::YFP; Bergs et al. under revision) using primers 5’-CCCCGGCGCGCCATCCATGAGCAGCATCACC-3’ and 5’-CCCCGAATTCCTTACTTGTACAGCTCGTCCAT-3’, and subcloned into vector XW12 with *Asc*I and *Eco*RI. Plasmid pOT17 (pQUAS::Δpes-10::PH-miniSOG(Q103L)) was generated as follows: PH-miniSOG(Q103L) sequence was amplified from plasmid pCZGY2849 (a gift from Andrew Chisholm: Addgene plasmid 74112) using primers 5’-CCCCGGCGCGCCCTTCGGATCCAGATCTATGCAC-3’ and 5’-TGTACAAGAAAGCTGGGTCG-3’) and subcloned into vector XW12 using *Asc*I and *Eco*RI restriction sites. The construct details are available on request.

### Animal tracking and behavioral analysis

For worms moving freely on NGM, locomotion parameters were acquired with a previously described worm tracker (Stirman et al., 2011) allowing to precisely target illumination of identified segments of the worm body by a modified off-the-shelf liquid crystal display (LCD) projector, integrated with an inverted epifluorescence microscope. Light power was measured with a powermeter (PM100, Thorlabs, Newton, NY, USA) at the specimen focal plane. Animals used in all the optogenetics experiments were raised in the dark at 20°C on NGM plates with *E. coli* OP50–1 and all-*trans*-retinal. The OP50-retinal plates were prepared 1–2 days in advance by seeding a 6 cm NGM-agar plate with 250 µl of OP50 culture and 0.25 µl of 100 mM retinal dissolved in ethanol. Young adults were transferred individually on plain NGM plates under red light (>600 nm) in a dark room and kept for 5 minutes in the dark before transfer to the tracker.

For experiments with ChR2 depolarizing MNs (Figs. 1, 2, Supplementary Fig. S1), blue light of 470 nm and 1.8 mW/mm^2^ intensity was used with the following light protocol: 20 s ’dark’ (referring to no blue light illumination) control, 20 s of illumination, followed again by 20 s dark. The animals’ body was divided into 11 segments, of which 3–10, 3–4, 5–6 or 9–10 were illuminated, depending on the experiment.

For optogenetic ablation experiments (Fig. 3A, Supplementary Fig. S2), AS MNs were ablated in animals expressing PH-miniSOG by 2.5 min exposure to 470 nm light of 1.8 mW/mm^2^ intensity; segments 3–10 out of 11 were illuminated. Animals were analyzed after a 2 h resting period for 60 s without illumination. Wild type worms were used as a control with the same illumination protocol. Ablation was verified by fluorescence microscopy in strain ZX2110 expressing green fluorescent protein (GFP) in all cholinergic neurons, in addition to PH-miniSOG in AS MNs.

For experiments of AS MN hyperpolarization using the histamine-gated Cl^-^-channel HisCl1 (Fig. 3B, Supplementary Fig. S2), worm locomotion was measured on NGM plates with 10 mM histamine 4 minutes after transfer from plates without histamine, for 60 s without illumination. The same strain on NGM without histamine served as a control.

For experiments in which MNs were hyperpolarized with natural Cl^-^-conducting anion channel rhodopsin (ACR1; Fig. 4), due to the high operational light sensitivity of the channel, the system was modified as described (Steuer Costa et al., 2017). An additional band pass filter (650 ± 25 nm) was inserted in the background light path and a mechanical shutter (Sutter Instrument Company, Novato, USA), synchronized to the light stimulation, was placed between projector and microscope. Control animals were tested for the background light stimulation and showed no response. The light stimulation protocol was 20 s without illumination, 20 s in 70 µW/mm^2^ 470 nm light and 20 s without illumination. The worms’ body was divided into 11 segments, and segments 3–10, 3–4, 5–6 or 9–10 were illuminated, respectively. As the experiment in *unc-47(e307)* background was performed with a different transgene injected into *unc-47(e307)* mutants, we tested the extrachromosomal array after outcrossing into wild type background, where it evoked contraction of the animals, as expected (Supplementary Fig. S3C).

Tracks were automatically filtered to exclude data points from erroneously evaluated movie frames with a custom-made workflow in KNIME (KNIME Desktop version 3.5, KNIME.com AG, Zurich, Switzerland; (Warr, 2012). Our constraints were that animals do not move faster than 1.25 mm/s and their length does not show a discrepancy above 25 % to the mean first five seconds of the video. Videos were excluded from analysis when more than 15 % of the data points had to be discarded by our constraints. Behavior data passed the Shapiro-Wilk normality test.

For determination of the ratio of dorso-/ventral angles (Fig. 1BIII, IV), the second out of 11 three point angles, measured from head to tail, was registered for animals for which the vulva position was previously indicated by manually indicating this to the software. For each track, values of the second three-point angle were averaged for dorsal and ventral bends individually, and the ratio was calculated.

To calculate the frame-to-frame difference of bending angles (Fig. 4E), data on each of eleven 3-point angles were extracted, smoothed by running an average of 15 frames, and the ∆ of absolute values between two subsequent frames were calculated and averaged for before and during illumination conditions for each angle. The light – no light ∆∆ of bending angles (Fig. 4F) were calculated by subtracting the value of the no light from the light condition. They were then averaged for the bending angles 1–5 (anterior) and 6–11 (posterior).

### Body posture analysis

Binarized videos of freely crawling animals were used to segment the animals’ body, and analyzed as described earlier (Hums et al., 2016; Stephens et al., 2008), using a custom MATLAB script (MathWorks, Natick, Massachusetts). Briefly, grey scale worm images were binarized with a global image threshold using Otsu’s method (Otsu, 1979). Objects encompassing border pixels were ignored and only the largest object was assumed to be the worm. The binary image was further processed (by thickening, removing spur pixels, flipping pixels by majority and filling holes). Worm skeletonization was achieved by thinning to produce an ordered vector of 100 body points and corresponding tangent angles (theta) from head to tail. Images that could not be analyzed or where the skeleton of the animal was unusually small were considered as missing data points. Head and tail assignment was checked manually. The theta angles were smoothened by a simple moving average with a window of 15 centered data points. The mean of these angles was then compared to the Eigenworms computed from previously published data on N2 videos (Stephens et al., 2008). The Eigen projections obtained were taken as a measure of worm posture and plotted.

### Electrophysiology

Recordings from dissected dorsal BWM cells of strain ZX2221 (used to avoid unspecific excitation of AS MNs via PINs were conducted as described previously (Liewald et al., 2008). Only the left side of the worm was cut to preserve commissural connections from the ventral nerve cord where AS MN cell bodies reside. Light activation was performed using an LED lamp (KSL-70, Rapp OptoElektronik, Hamburg, Germany; 470 nm, 8 mW/mm^2^) and controlled by an EPC10 amplifier and Patchmaster software (HEKA, Germany).

### Ca^2+^ imaging microscope setup

Fluorescence measurements were carried out on an inverted fluorescence microscope (Axiovert 200, Zeiss, Germany) equipped with motorized stage MS 2000 (Applied Scientific Instrumentation, USA) and the PhotoTrack quadrant photomultiplier tube (PMT; Applied Scientific Instrumentation, USA). Two high-power light emitting diodes (LEDs; 470 and 590 nm wavelength, KSL 70, Rapp Optoelektronik, Germany) or a 100 W HBO mercury lamp were used as light sources. A Photometrics DualView-Λ beam splitter was used to obtain simultaneous dual-wavelength acquisition; these were coupled to a Hamamatsu Orca Flash 4.0 sCMOS camera operated by HCImage Live (Hamamatsu) or MicroManager (http://micro-manager.org). Light illumination protocols (temporal sequences) were programmed on, and generated by, a Lambda SC Smart shutter controller unit (Sutter Instruments, USA), using its TTL output to drive the LED power supply or to open a shutter when using the HBO lamp.

### Measurement of Ca^2+^ in muscles and AS MNs in immobilized worms

For measurements of GCaMP3 (Fig. 2) and RCaMP (Fig. 3B) in muscles and GCaMP6 in AS MNs (Figs. 5, 6), the following light settings were used: GFP/mCherry Dualband ET Filterset (F56–019, AHF Analysentechnik, Germany), was combined with 532/18 nm and 625/15 nm emission filters and a 565 longpass beamsplitter (F39–833, F39–624 and F48–567, respectively, all AHF). ChR2 stimulation was performed using 1.0–1.2 mW/mm^2^ blue light, unless otherwise stated. To measure RCaMP or mCherry fluorescence, 590 nm, 0.6 mW/mm^2^ yellow light was used. The 2x binned images were acquired at 50 ms exposure time and 20 fps. Animals were immobilized on 2 or 4% M9 agar pads with polystyrene beads (Polysciences, USA) and imaged by means of 25x or 40x oil objective lenses. 5 s of yellow light illumination and 15 s of blue light illumination protocols were used. For RCaMP imaging 20 s yellow light illuminations were used. Measurements of control animals (i.e. raised without ATR, or without histamine) were conducted the same way as for animals kept in the presence of ATR, or exposed to histamine.

Image analysis was performed in ImageJ (NIH). For Ca^2+^-imaging in muscles, regions of interest (ROIs) were selected for half of the BWM cells in the field of view, or around the neuron of interest for Ca^2+^-imaging in AS MNs. Separate ROIs were selected for background fluorescence with the same size. Mean intensity values for each video frame were obtained and background fluorescence values were subtracted from the fluorescence values derived for GCaMP or RCaMP. Subtracted data was normalized to ΔF/F = (F_i_-F)/F, where F_i_ represents the intensity at the given time point and F represents the average fluorescence of the entire trace.

### Measurement of Ca^2+^ in muscles and AS motor neurons in moving animals

Measurements of GCaMP6 and mCherry were performed using the same filter and microscope settings as for immobilized worms. Moving worms were assayed on 1% agar pads in M9 buffer. Tracking was based on the PhotoTrack system (Applied Scientific Instrumentation, USA) that uses the signals from a 4-quadrant photomultiplier tube (PMT) sensor for automated repositioning of a motorized XY stage to keep a moving fluorescent marker signal in the field of view (Faumont et al., 2011). For this purpose, an oblique 80% transmission filter was inserted in the light path to divert 20% of the light to the PMT quadrants. A 535/30 bandpass filter (F47–535, AHF) was used to narrow the emission spectrum prior to detection for improved tracking performance. A fluorescent marker GFP was expressed in vulval muscle cells, strains PD4665 and ZX2012. Video files containing data of both fluorescent channels (for GCaMP6 and mCherry) were processed with custom written Wolfram Mathematica notebooks. Both color channels were virtually overlaid to accurately correct the spatial alignment. Images were first binarized to identify the centroid of the moving neuronal cell bodies throughout all frames. Mean intensity values of a circular ROI (18 pixel radius) centered on this centroid were measured and subtracted with the mean intensity values of a surrounding donut shaped background ROI (5 pixel width). Coordinates of two AS neurons of interest were recorded relative to the vulva and to each other to obtain their relative distance and the angle between the vulva and the two neurons of interest. The traces were normalized to ΔF/F = (F_i_-F)/F, where F represents the average of the entire trace, and were used for correlation analysis.

### Measurement of Ca^2+^ signals in PINs

Ca^2+^ imaging with cameleon D3cpv (Palmer et al., 2006) was performed on 5% agar pads as described (Kawano et al., 2011) on an Axiovert 200 microscope (Zeiss), using a 100x/1.30 EC Plan-Neofluar Oil M27 oil immersion objective. ChR2 stimulation was performed using 8 mW/mm^2^ blue light delivered by a 100 W HBO mercury lamp. The excitation light path was split using a dual-view (Photometrics) beam splitter with a CFP/YFP filter set. The YFP/CFP ratio after background subtraction was normalized to the ΔR/R=(R_i_-R)/R, where R_i_ represents the YFP/CFP ratio at the given time point and R represents the average of the entire trace during blue light stimulation. YFP/CFP ratios without normalization were used for quantification and statistics (Figure 6B IV). This data did not pass the Shapiro-Wilk normality test.

### Correlation analysis

Cross-correlation analyses were performed with built-in MATLAB functions. Ca^2+^ transients in AS6 and AS7 and vulva angles were smoothed for 10 frames. Individual bending events identified as segment of the trace between two minima were used for cross-correlation with 100 time lags (10 s). For comparison of peak correlations, the maximum correlation (positive or negative) in a 5 s time window centered on the peak of the control mean correlation was used.

### Statistics

Data is given as means ± SEM. Significance between data sets after two-tailed Student’s t-test or after Mann-Whitney U-test or after ANOVA is given as p-value (* p ≤0.05; ** p ≤ 0.01; *** p ≤ 0.001), the latter was given following Tukey’s post-hoc test. Data was analyzed and plotted in Excel (Microsoft, USA), in OriginPro 2016 (OriginLab Corporation, Northampton, USA) or in MATLAB (MathWorks, Natick, Massachusetts, USA).

## AUTHOR CONTRIBUTIONS

O.T. designed experiments, performed experiments, analyzed data, wrote the manuscript; P.V.d.A. performed experiments and provided analysis code; A.B. provided plasmids. T. G. and O. B. generated plasmids and strains and performed initial experiments. W.S.C provided analysis code and contributed to data analysis. J.F.L. performed experiments and analyzed data; A.G. supervised the project, designed experiments, performed data interpretation, and edited the manuscript.

## ACKNOWLEDGEMENTS

We thank Cori Bargmann for suggesting to study the AS MNs. We are grateful to Mei Zhen, Kang Shen, Xing Wei, Navin Pokala, Cori Bargmann, Yishi Jin, and Andrew Chisholm for reagents and to Isabell Franz, Mona Hoeret, Heike Fettermann, Regina Wagner and Heinz Schewe for expert technical assistance. Yongmin Cho, Daniel Porto and Hang Lu provided equipment and software. We thank Gal Haspel for fruitful discussions. This work was funded by a GO-IN stipend of Goethe University, in conjunction with the EU program PCOFUND-GA-2011–291776, GO-IN (to O.T.), by a IMPReS PhD stipend (to A.B.) and by grants GO1011/4–2 (Protein-based Photoswitches), GO1011/8–1 (NewOptogeneticsTools) and EXC115/3 (Cluster of Excellence Frankfurt - Macromolecular Complexes) from the Deutsche Forschungsgemeinschaft (DFG) to AG.

## Supplementary Figures

**Figure S1. AS MN photostimulation induces postsynaptic currents and local AS neuron activation affects body length: A)** Representative postsynaptic currents evoked in dorsal muscle cell in response to photodepolarization (indicated by blue bar) of the AS MNs via ChR2. **B)** I, II) Speed and III, IV) body length (time traces, I-IV, and group data, V, VI) before and during photodepolarization of AS MNs or of all cholinergic MNs in the anterior, middle and posterior segments of the worm body by ChR2 (in animals raised with ATR). P value * ≤ 0.05, ** ≤ 0.01, *** ≤ 0.001; number of animals is indicated. Statistical test: ANOVA with Tukey’s post-hoc test.

**Figure S2. Optogenetic inactivation and HisCl1-induced hyperpolarization of AS MNs affects locomotion speed and bending angles: A)** Time traces of mean (± SEM) speed (I) and bending angles (II) of freely moving wild type animals, or animals expressing PH-miniSOG in AS MNs, 2 h after 150 s of blue light exposure. **B)** Time traces of mean (± SEM) speed of freely moving animals expressing HisCl1in AS MNs, comparing animals on plates without and with 10 mM histamine. Number of animals tested is indicated.

**Figure S3. Local stimulation of AS MNs in different body segments: A)** Schematic showing how the 13 points defining 11 3-point angles correspond to the 11 body segments that were individually illuminated. **B)** Mean speed (I. II) and body length (III, IV) of animals expressing ACR1 in AS MNs, in which the indicated body segments were illuminated. Group data shown in II, IV. **C)** The extrachromosomal array expressing ACR1 in the *unc-47(e307)* mutant as shown in main Fig. 4B, C functions as expected in wt background. P value * ≤ 0.05; number of animals is indicated. Statistical test in B: ANOVA with Tukey’s post-hoc test.

**Figure S4. AS MNs are simultaneously activated by photostimulation of the AVA and AVB PINs: A, B)** Cross-correlation analysis of GCaMP6 fluorescence signals (ΔF/F) in AS3 and AS8 neurons, during photodepolarization of AVA (A) or AVB (B), expressing ChR2, respectively, in animals raised in the presence of ATR.

## Supplementary Videos

**Video 1.** Freely moving animal before and during photodepolarization of AS MNs by ChR2 (in animal raised with ATR), blue light = 470 nm, 1.8 mW/mm^2^. Video plays at 3x speed (75 f/s).

**Video 2.** Ca^2+^ signal in the BWM expressing GCaMP3 during photodepolarization of AS MNs by ChR2 (in animal raised with ATR), blue light = 470 nm, 1.2 mW/mm^2^. Video plays at 3x speed (75 f/s).

**Video 3.** Freely moving animal, 2 hours after optogenetic ablation (150 s) of AS MNs by PH-miniSOG.

**Video 4.** Freely moving animal expressing HisCl1 after 4 min exposure on 10 mM histamine. Video plays at 3x speed (75 f/s).

**Video 5.** Freely moving animal expressing ACR1 in AS MNs before (20 s) and during ACR1 photoactivation by 470 nm 1.8 mW/mm^2^ blue light. Video plays at 3x speed (75 f/s).

**Video 6.** Selective illumination of anterior segment. Freely moving animal expressing ACR1 in AS MNs before (20 s) and during ACR1 photoactivation by 470 nm 1.8 mW/mm^2^ blue light. Video plays at 3x speed (75 f/s).

**Video 7.** Selective illumination of midbody segment. Freely moving animal expressing ACR1 in AS MNs before (20 s) and during ACR1 photoactivation by 470 nm 1.8 mW/mm2 blue light. Video plays at 3x speed (75 f/s).

**Video 8.** Selective illumination of posterior segment. Freely moving animal expressing ACR1 in AS MNs before (20 s) and during ACR1 photoactivation by 470 nm 1.8 mW/mm^2^ blue light. Video plays at 3x speed (75 f/s).

**Video 9.** Moving animal expressing GCaMP6 and mCherry in the AS MNs while the animal is being automatically tracked *via* the GFP marker in vulva muscles. Video plays at 0.35x speed (7 f/s).

**Video 10.** Ca^2+^ signal in the AS MNs expressing GCaMP6 during photodepolarization of the AVA PIN by ChR2 (in animal raised with ATR), 470 nm blue light, 1.2 mW/mm^2^. Video plays at 3x speed (75 f/s).

**Video 11.** Ca^2+^signal in the AS MNs expressing GCaMP6 during photodepolarization of AVB by ChR2 (in animal raised with ATR), 470 nm blue light, 1.2 mW/mm^2^. Video plays at 3x speed (75 f/s).

